# Standard Flow Multiplexed Proteomics (SFloMPro) – An Accessible and Cost-Effective Alternative to NanoLC Workflows

**DOI:** 10.1101/2020.02.25.964379

**Authors:** Conor Jenkins, Ben Orsburn

## Abstract

Multiplexed proteomics using isobaric tagging allows for simultaneously comparing the proteomes of multiple samples. In this technique, digested peptides from each sample are labeled with a chemical tag prior to pooling sample for LC-MS/MS with nanoflow chromatography (NanoLC). The isobaric nature of the tag prevents deconvolution of samples until fragmentation liberates the isotopically labeled reporter ions. To ensure efficient peptide labeling, large concentrations of labeling reagents are included in the reagent kits to allow scientists to use high ratios of chemical label per peptide. The increasing speed and sensitivity of mass spectrometers has reduced the peptide concentration required for analysis, leading to most of the label or labeled sample to be discarded. In conjunction, improvements in the speed of sample loading, reliable pump pressure, and stable gradient construction of analytical flow HPLCs has continued to improve the sample delivery process to the mass spectrometer. In this study we describe a method for performing multiplexed proteomics without the use of NanoLC by using offline fractionation of labeled peptides followed by rapid “standard flow” HPLC gradient LC-MS/MS. Standard Flow Multiplexed Proteomics (SFloMPro) enables high coverage quantitative proteomics of up to 16 mammalian samples in about 24 hours. In this study, we compare NanoLC and SFloMPro analysis of fractionated samples. Our results demonstrate that comparable data is obtained by injecting 20 μg of labeled peptides per fraction with SFloMPro, compared to 1 μg per fraction with NanoLC. We conclude that, for experiments where protein concentration is not strictly limited, SFloMPro is a competitive approach to traditional NanoLC workflows with improved up-time, reliability and at a lower relative cost per sample.

Data are available via ProteomeXchange with identifier PXD016704.

Abstract Graphic

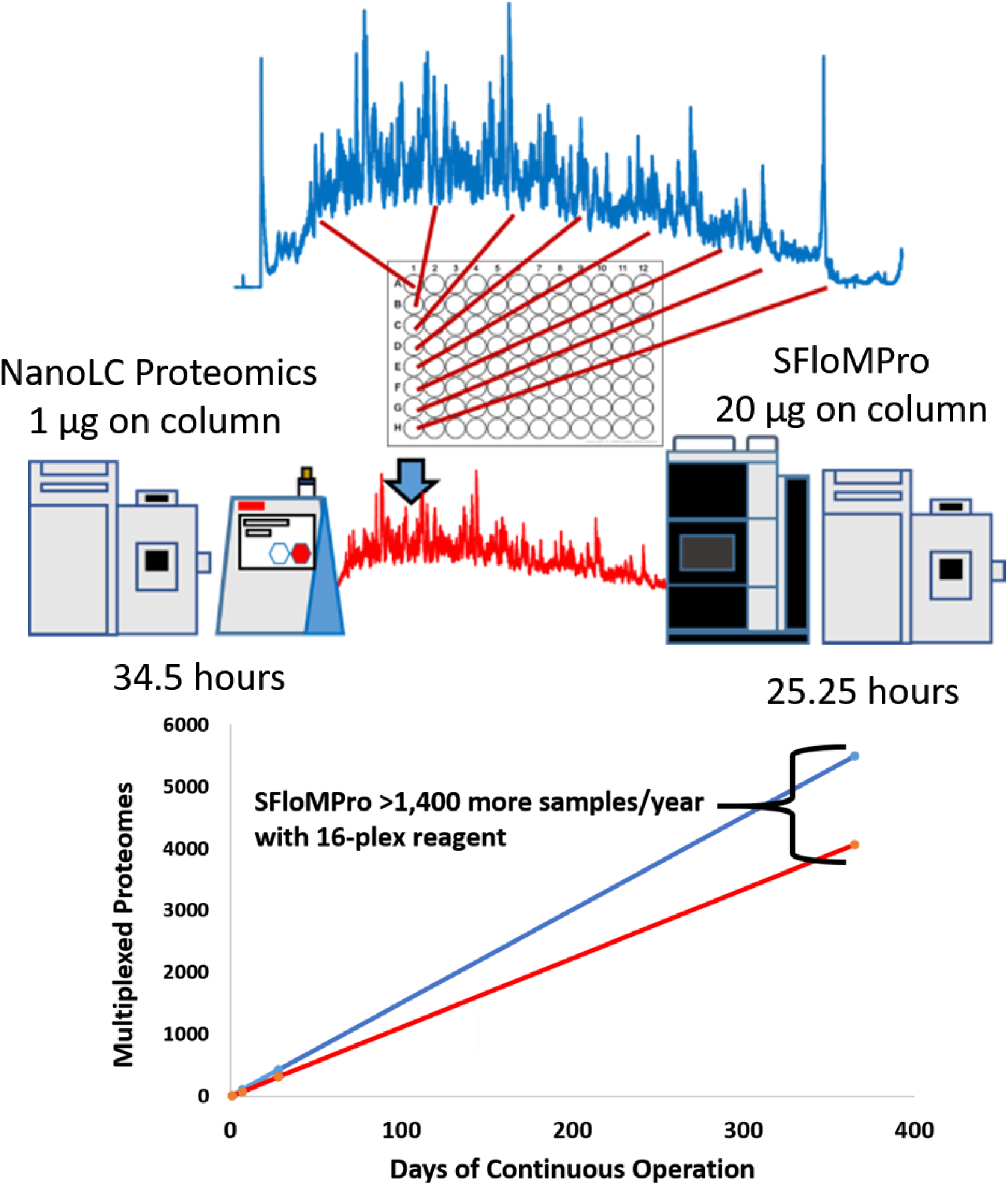

## Introduction

Shotgun proteomics has several technical challenges that currently inhibits its widespread adoption. Many of these have been addressed by recent advances in mass spectrometry engineering leading to marked increases in accuracy, resolution, speed and sensitivity. Both instrument data acquisition and post processing software have improved, with the transition of modern algorithms into graphic user interfaces (GUIs)^1234^

A major challenge in shotgun proteomics is the use of nanoflow liquid chromatography (NanoLC). In LC-MS based shotgun proteomics samples are diluted by chromatography running buffer. Due to the relatively low sensitivity of the first tandem mass spectrometers utilized for global proteomics, concentrating the sample with the use of progressively lower flow rates was an attractive solution.^5^ The resulting methodology, NanoLC, was rapidly adopted in shotgun proteomics, to the point that it has been recently described as dogma.^6^

NanoLC requires the precise plumbing of fragile fused silica columns with internal diameters ranging 20 - 100 μm and flow rates typically ranging 20 - 500 nL per minute. As few pumps existed that could reliably deliver these flow rates, early practitioners utilized solvent splits from higher flow pumps that would divert as much as 99% of the utilized solvent to waste with the remaining 1% used for peptide gradient elution.^7^

Recent advances have seen the utilization of pressurized gas-driven, low capacity syringe pumps that can directly construct reversed phase chromatography gradients of these flow rates without solvent splits. Furthermore, the fused silica lines are now often coated in plastic sheaths and integrate “zero dead volume” union connections that simplify the plumbing of NanoLC systems.

In order to work with nL/min flow rates, even modern systems require more maintenance to prevent and remove the presence of trapped atmospheric air and evaporated solvents when compared to conventional flow technologies. The user manual for the NanoLC system recommends daily cycles of solvent purges and programs to flush air from pumps (Supplemental Figure 1). In contrast, an ultra-high pressure (UHPLC) system purchased at this same date from the same vendor recommends solvent flushes at monthly intervals (Supplemental Figure 1). Despite this increase in maintenance, the NanoLC is orders of less reproducible than higher flow LC systems. UHPLC conventional flow systems have shown peak retention times varying less than one second over hundreds of samples.^1^ It is well accepted that NanoLC experiments have less reproducible retention times.^8^ The recently described IonStar system for clinical proteomics carefully controls all chromatography variables for maximum NanoLC reproducibility using 100 cm columns, but still requires adjustments to align retention times that differ by as much as 60 seconds between runs.^9^

Common quantitative proteomic experiments utilize isobaric tags such as the commercial iTRAQ and TMT products.^10^ By performing relative quantification in the MS/MS spectra of tagged peptides, peptide retention times and NanoLC retention time reproducibility are less of a concern. In this procedure, peptides from independent samples are each labeled with a unique isobaric tag and then combined for LCMS. The relative abundances of the peptides are revealed in MS/MS or MS/MS/MS fragmentation spectra and sample to sample quantitation can be achieved. A critical step in the experiment to minimize bias is the efficiency of the peptide labeling. In order to obtain complete labeling of the peptides, the manufacturer of the reagents recommends a ratio of 1:8 peptide to label.

However, recent studies have demonstrated effective complete labeling of peptides with a lower ratio of peptides, with 1:4 and even 1:2 shown to be effective.^11,12^ Currently, the smallest aliquot commercially available for each labeling compound is 800 μg of label per sample. This appears to constitute a large discrepancy, as the typical upper limit for peptides for NanoLC separation, is considered 0.2 – 4 μg of peptide injection.^13^ This results in the generation of excess materials in either a large amount of unused label or labeled peptide. In addition, due to the high reactivity of the label, these excess materials are often considered unsuitable for use following rehydration and are typically discarded as waste.^11^ Alternative approaches to NanoLC are gaining in popularity, with multiple studies using capillary zone electrophoresis^14,15^ and microflow chromatography^16,17^demonstrating promise as alternative methodologies.

Recent work has also described the use of “standard flow” proteomics, reaching the conclusion that flow rates in the 50 - 200 μl per minute range can provide quality proteomics data with proper optimization.^6,18^ Building on this work, we describe a standard flow multiplexed proteomics (SFloMPro) workflow. Our results show that SFloMPro produces comparable data to that of NanoLC but requires 20 times more labeled peptide on column. Due to the rapid preparation of LC gradients and sample loading of conventional flow uHPLC systems, we demonstrate a time savings of nearly 25% with no changes in our sample preparation workflow or reagent usage. By removing the requirements of the purchase of a NanoLC system and the technical hurdles associated with this technology, SFloMPro requires only a HPLC-Orbitrap instrument configuration to perform quantitative proteomics.

Using SFloMPro, we performed multiplexed quantification of over 8,000 mammalian proteins in approximately 24 hours of total instrument acquisition time. We conclude that SFloMPro is an accessible alternative for high throughput proteomics in conditions such as cell culture or any biological samples where sample abundance is not a strictly limiting factor.

## Materials and Methods

### Cells and cell culture

One T25 flask of murine cell culture line BalbC was prepared per condition. Cells were harvested lysed, resulting in 1-4 mg total protein per condition, as quantified by BCA. Approximately 200 μg of protein was utilized for digestion with reduction and alkylation of the cysteines with DTT and iodoacetamide, respectively. Six channels were labeled with TMT 11-plex reagent in a 1:4 ratio of peptide to label. The following channels were utilized in this study: 129N,129C, 130N, 130C, 131N, 131C, where the last channel is pooled samples.

### High pH reversed phase fractionation

The combined labeled peptides were separated on a Thermo Accela 1250 HPLC using a Waters XBridge BEH130 C18 3.5μm 2.1mm × 150 mm column using a flow rate of 0.2 mL/min with Buffer A and B as 25mM Ammonium bicarbonate pH 8.0 in LC-MS grade water and LC-MS grade acetonitrile (Fisher), respectively. Peptides were separated on a linear gradient of 5 - 35% B over 60 min, with a linear increase to 70% B over 12 min. Fractions were collected every 45 sec using a Foxy Junior fraction collector to result in complete filing of a 96 well plate. Stepwise concatenation was performed by incrementally combining every 24^th^ well, for example: sample 1 is combined results of wells 1, 25, 49, and 73 combined; sample 2 is combined result of wells 2, 26, 50, and 74. The 24 resulting fractions were desalted with Pierce spin columns (part number 89851) and SpeedVac to near dryness for LC-MS analysis.

### NanoLC Separation

Approximately 1 μg of peptides were loaded by EasyNLC 1000 (Thermo Fisher) onto a 3 mm PepMap desalting column prior to solvent loading gradient elution on the 15 cm EasySpray 3 μm PepMap column. Buffers consisted of 0.1% formic acid in LCMS grade water (A) and 80% LCMS grade acetonitrile (B) from Thermo. Prior to each injection the pre-column was equilibrated with 1 μL Buffer A at a maximum pressure of 600 bar. The analytical column was equilibrated the same with 6 μL Buffer A.

For each injection, 2 μL of sample was picked up and a total of 6 μL of sample plus loop loading buffer was loaded to the trap column prior to closing the waste valve and beginning the gradient. The gradient consisted of a two-stage gradient with a condition of 5% B at 300 nL/min that increased to 24% B in 31 min, followed by an increase to 38% B by 56 minutes. The column was reconditioned by increasing the flow rate to 500 nL/min and ramping to 98% B by 65 min before column equilibrations for the next sample.

### uHPLC Separation and ionization conditions

All analyses used a Vanquish H uHPLC (Thermo) coupled to Q Exactive mass spectrometer. Starting conditions were based on a recent report by Lenco *et al*.^6^ using a Waters BEH Peptide BEH C18 1.7μm × 2.1 × 150mm column. Optimization of injection and gradient was performed using the HeLa peptide standard (Pierce). The final gradient utilized 5% DMSO and 0.1% formic acid with Milli-Q water as Buffer A and HPLC grade acetonitrile as Buffer B. The gradient began at 200 μL/min 0% B and ramped to 5% B in 5 min followed by an increase to 30% B by 50 min, an increase to 90% B in 6 min with increase to 0.4 mL/min for the remainder of the run, with a 1 min hold before resuming to baseline conditions to a total run length of 65 min. All gradient changes were with a pump curve of 5 arbitrary units. The Q Exactive Classic system was equipped with HESI-II system (Thermo) and used the following source conditions described by the Xcalibur Tune software for 0.2 mL/min flow rates: ESI voltage 3500, capillary temperature 325, sheath gas 45, auxiliary gas 10, spare gas 2, probe heater temperature of 150C and S-lens RF of 70.

### Mass spectrometer conditions

An identical data-dependent acquisition method was used for both experiments. MS1 spectra were acquired at a resolution of 70,000 with an AGC target of 3×10^6^ charges or 100 ms maximum ion injection time. MS1 was acquired 381 - 1581 m/z. The top 10 ions were selected for MS/MS using a resolution of 35,000 at m/z of 200 with an AGC target of 1 ×10^6^ or maximum ion injection time of 114 ms. An isolation window of 2.2 Da was used along with a fixed first mass of 110 m/z. A normalized collision energy of 30 was used for both experiments. Ions of unassigned, single charge, or charge state of 8 or above were excluded from MS/MS and dynamic exclusion was used with a 30 sec window.

### Data Processing

All data was processed in Proteome Discoverer 2.4 (Thermo). For TMT quantitative analysis the manufacturer default workflow for reporter ion quantification for TMT 11-plex reagent. The SeQuestHT search engine was used with the following settings: 10ppm MS1 tolerance, 0.02 Da MS/MS tolerance, variable modifications of methionine oxidation and acetylation of the protein N terminus, along with static modifications of carbamidomethylation of cysteines and the addition of the TMT 6/10/11 plex tag on peptide N-terminus and lysine. Up to two missed cleavage events were allowed. To evaluate relative peak widths and other variables the samples were reran as described, with the removal of the reporter ion quantification node and replacement with the Minora feature alignment and peak detection node using default parameters.

## Results and Discussion

### Total Instrument Acquisition Time

In the increasingly competitive research core environment the number of samples completed per unit time may be a critical measurement for financial success.^19^ A recent study that utilized high levels of high pH offline and rapid NanoLC gradients noted the minimum loading time of 22 min for a nanoflow LC system similar to the one employed in this study.^20^ In an attempt to keep variable consistent the total acquisition time for the NanoLC and SFloMPro samples were set at approximately 60 minutes. However, the time stamps embedded in the manufacturer binary files indicates that the 24 NanoLC samples required 33.5 hours from first to final sample. In contrast, the 24 SFloMPro samples were only separated by 25.3 hours, a 25% reduction in run time. While multiple approaches have been demonstrated for parallel trapping and elution in NanoLC proteomics, the systems in this lab do not have these capabilities.^21,22^

### Peptide and protein identifications

Table 1 is a summary of the two experiments. Full results are available in Supplemental Data I. Although the NanoLC experiment acquired more total MS/MS spectra and peptide spectral matches (PSMs), this did not translate to a corresponding increase in the number of peptides and proteins identified. An average of 2.1 PSMs were identified for each peptide in the NanoLC files, compared to 1.4 PSMs in the SFloMPro file set. The mass spectrometry proteomics data have been deposited to the ProteomeXchange Consortium via the PRIDE partner repository with the dataset identifier PXD016704 and 10.6019/PXD016704.

**Table 1.** An overview of the results of the two experiments described.

| Experiment | MS/MS spectra | PSMs | Peptides | Proteins | Total Run Time (hr) |
| --- | --- | --- | --- | --- | --- |
| NanoLC | 488,715 | 137,460 | 65,295 | 7,685 | 34.50 |
| SFloMPro | 359,253 | 107,552 | 74,160 | 8,086 | 25.25 |

For analysis of relative chromatography conditions, the two datasets were reprocessed using the Minora Feature Detector node. One output of the node is the identification of left and right retention times for each chromatographic feature. From these measurements, the average peak widths for the NanoLC experiment was determined to be 48.6 ± 7.2 sec. The uHPLC system recorded a peak width of 34.8 ± 0.74 sec. The use of the identical dynamic exclusion settings of 30 sec appears to have been suboptimal for the NanoLC experiment and corresponds to repeated fragmentation of the same peptides, thereby reducing peptide and total protein identifications. This observation suggests that further optimization of the NanoLC and MS/MS conditions are appropriate due to the wider relative peaks in NanoLC compared to uHPLC using the resources described here.

As shown in Figure 1 the overlap in protein group identifications between the two methods is in high concordance, with proteins greater than 2 peptides per protein shown in the graph. Considering the well-recognized variability in protein group identifications due to parsimony, these results likely approach the expected theoretical maximum when comparing two separate experiments.^23^ Furthermore, Figure 2 is a demonstration of the normalized loading plots of each channel. Box plots of the same color represent the total normalized loading of the same channel compared between the NanoLC and SFloMPro experiments and further demonstrates the relative level of concordance between the two sample sets.

**Figure 1.**
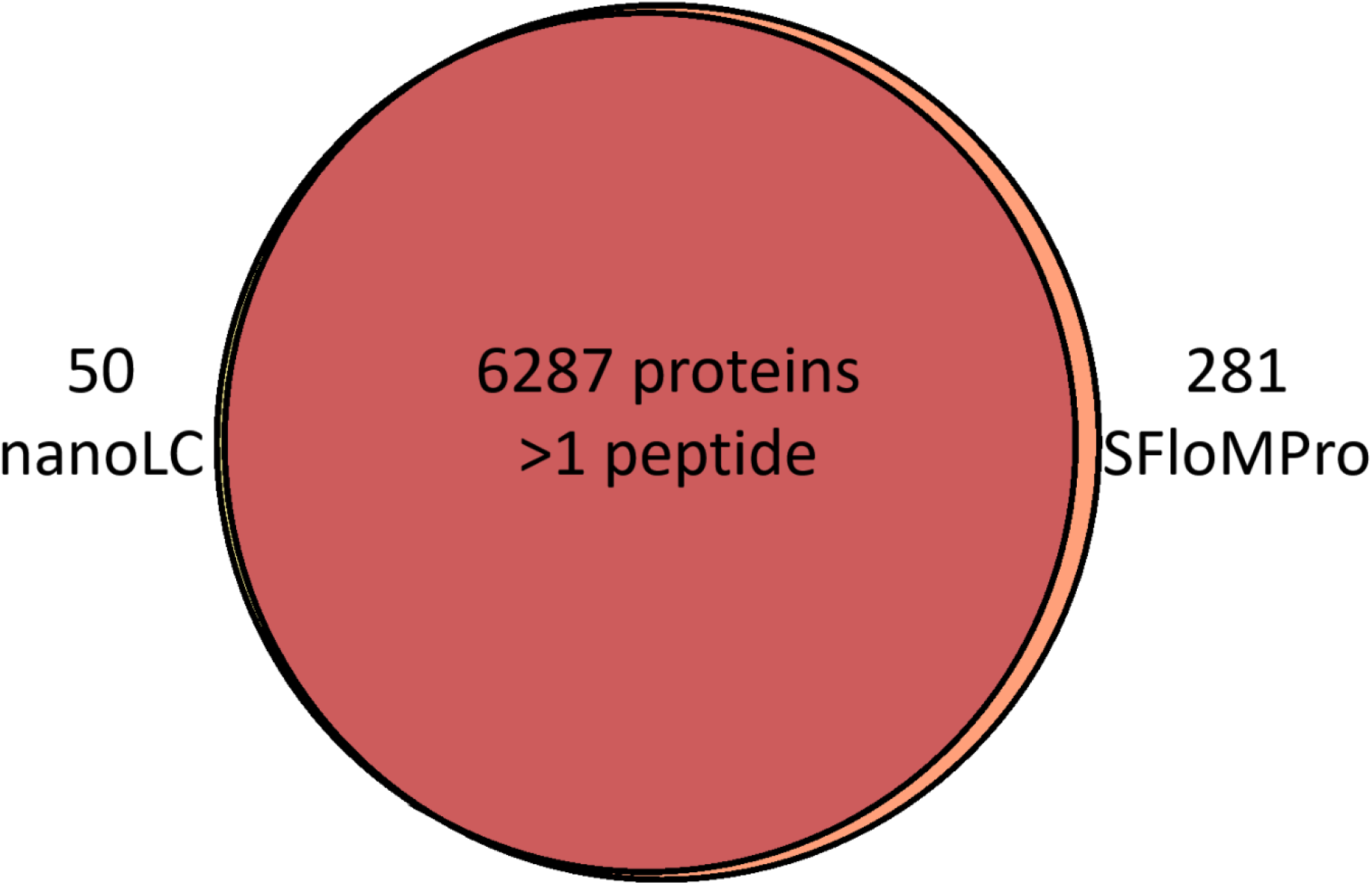
A scaled Venn diagram showing the overlap in protein identifications when two or more unique peptides are used as a filter.

**Figure 2.**
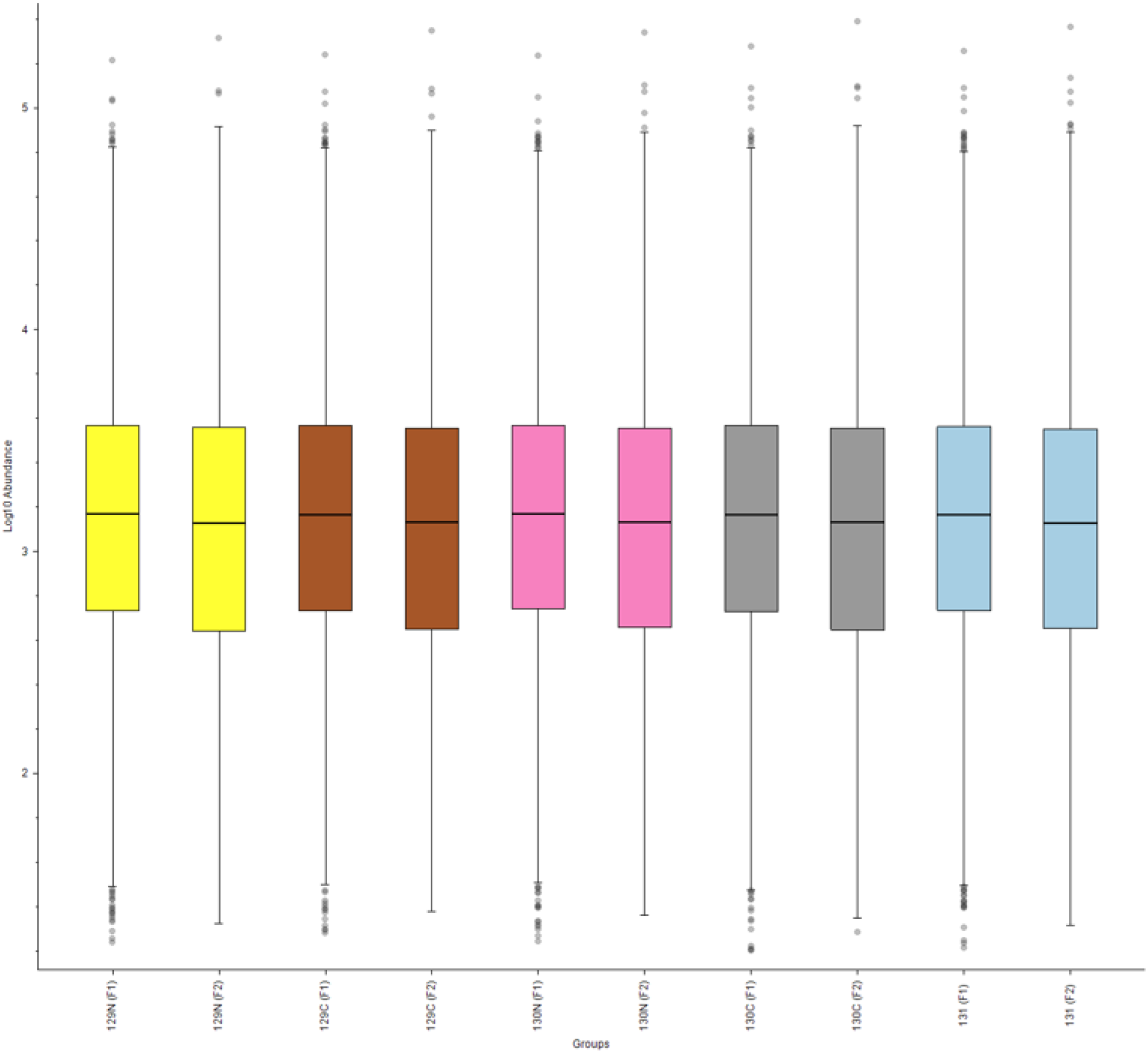
Normalized loading plots for each channel utilized. Matching colors are the same TMT channel. F1 represents the NanoLC experiment while F2 represents the SFloMPro samples.

In this study, we describe the SFloMPro shotgun proteomics workflow which circumvents technical challenges in the field. By eliminating the NanoLC from the experiment and optimizing analytical flow HPLC, we obtain nearly equivalent data. Furthermore, we find that SFloMPro Proteomics is an economical alternative due increased speed in sample loading and gradient equilibration as well as to lower relative costs for consumables and service items.

## References

(1) Millikin, R. J.; Solntsev, S. K.; Shortreed, M. R.; Smith, L. M. Ultrafast Peptide Label-Free Quantification with FlashLFQ. J. Proteome Res. 2018. https://doi.org/10.1021/acs.jproteome.7b00608.

(2) Cox, J.; Neuhauser, N.; Michalski, A.; Scheltema, R. A.; Olsen, J. V.; Mann, M. Andromeda: A Peptide Search Engine Integrated into the MaxQuant Environment. J. Proteome Res. 2011. https://doi.org/10.1021/pr101065j.

(3) Weisser, H.; Nahnsen, S.; Grossmann, J.; Nilse, L.; Quandt, A.; Brauer, H.; Sturm, M.; Kenar, E.; Kohlbacher, O.; Aebersold, R.; et al. An Automated Pipeline for High-Throughput Label-Free Quantitative Proteomics. J. Proteome Res. 2013. https://doi.org/10.1021/pr300992u.

(4) Dorfer, V.; Pichler, P.; Stranzl, T.; Stadlmann, J.; Taus, T.; Winkler, S.; Mechtler, K. MS Amanda, a Universal Identification Algorithm Optimized for High Accuracy Tandem Mass Spectra. J. Proteome Res. 2014. https://doi.org/10.1021/pr500202e.

(5) Zhang, Y.; Fonslow, B. R.; Shan, B.; Baek, M. C.; Yates, J. R. Protein Analysis by Shotgun/Bottom-up Proteomics. Chemical Reviews. 2013. https://doi.org/10.1021/cr3003533.

(6) Lenčo, J.; Vajrychová, M.; Pimková, K.; Prokšová, M.; Benková, M.; Klimentová, J.; Tambor, V.; Soukup, O. Conventional-Flow Liquid Chromatography-Mass Spectrometry for Exploratory Bottom-Up Proteomic Analyses. Anal. Chem. 2018. https://doi.org/10.1021/acs.analchem.8b00525.

(7) Mellors, J. S.; Gorbounov, V.; Ramsey, R. S.; Ramsey, J. M. Fully Integrated Glass Microfluidic Device for Performing High-Efficiency Capillary Electrophoresis and Electrospray Ionization Mass Spectrometry. Anal. Chem. 2008. https://doi.org/10.1021/ac800428w.

(8) Lawrence, R. T.; Searle, B. C.; Llovet, A.; Villén, J. Plug-and-Play Analysis of the Human Phosphoproteome by Targeted High-Resolution Mass Spectrometry. Nat. Methods 2016. https://doi.org/10.1038/nmeth.3811.

(9) Shen, X.; Shen, S.; Li, J.; Hu, Q.; Nie, L.; Tu, C.; Wang, X.; Orsburn, B.; Wang, J.; Qu, J. An IonStar Experimental Strategy for MS1 Ion Current-Based Quantification Using Ultrahigh-Field Orbitrap: Reproducible, In-Depth, and Accurate Protein Measurement in Large Cohorts. J. Proteome Res. 2017. https://doi.org/10.1021/acs.jproteome.7b00061.

(10) Kammers, K.; Cole, R. N.; Tiengwe, C.; Ruczinski, I. Detecting Significant Changes in Protein Abundance. EuPA Open Proteom 2015, 7, 11–19. https://doi.org/10.1016/j.euprot.2015.02.002.

(11) Navarrete-Perea, J.; Yu, Q.; Gygi, S. P.; Paulo, J. A. Streamlined Tandem Mass Tag (SL-TMT) Protocol: An Efficient Strategy for Quantitative (Phospho)Proteome Profiling Using Tandem Mass Tag-Synchronous Precursor Selection-MS3. J. Proteome Res. 2018. https://doi.org/10.1021/acs.jproteome.8b00217.

(12) Zecha, J.; Satpathy, S.; Kanashova, T.; Avanessian, S. C.; Kane, M. H.; Clauser, K. R.; Mertins, P.; Carr, S. A.; Kuster, B. TMT Labeling for the Masses: A Robust and Cost-Efficient, in-Solution Labeling Approach. Mol. Cell. Proteomics 2019. https://doi.org/10.1074/mcp.TIR119.001385.

(13) Shen, S.; Wang, X.; Orsburn, B. C.; Qu, J. How Could IonStar Challenge the Current Status Quo of Quantitative Proteomics in Large Sample Cohorts? Expert Review of Proteomics. 2018. https://doi.org/10.1080/14789450.2018.1490646.

(14) Zhang, Z.; Hebert, A. S.; Westphall, M. S.; Qu, Y.; Coon, J. J.; Dovichi, N. J. Production of over 27 000 Peptide and Nearly 4400 Protein Identifications by Single-Shot Capillary-Zone Electrophoresis-Mass Spectrometry via Combination of a Very-Low-Electroosmosis Coated Capillary, a Third-Generation Electrokinetically-Pumped Sheath-Fl. Anal. Chem. 2018. https://doi.org/10.1021/acs.analchem.8b02991.

(15) Wojcik, R.; Li, Y.; MacCoss, M. J.; Dovichi, N. J. Capillary Electrophoresis with Orbitrap-Velos Mass Spectrometry Detection. Talanta 2012. https://doi.org/10.1016/j.talanta.2011.10.048.

(16) Vowinckel, J.; Zelezniak, A.; Bruderer, R.; Mülleder, M.; Reiter, L.; Ralser, M. Cost-Effective Generation of Precise Label-Free Quantitative Proteomes in High-Throughput by MicroLC and Data-Independent Acquisition. Sci. Rep. 2018. https://doi.org/10.1038/s41598-018-22610-4.

(17) Bian, Y.; Zheng, R.; Bayer, F. P.; Wong, C.; Chang, Y. C.; Meng, C.; Zolg, D. P.; Reinecke, M.; Zecha, J.; Wiechmann, S.; et al. Robust, Reproducible and Quantitative Analysis of Thousands of Proteomes by Micro-Flow LC–MS/MS. Nat. Commun. 2020. https://doi.org/10.1038/s41467-019-13973-x.

(18) Fernández-Niño, S. M. G.; Smith-Moritz, A. M.; Chan, L. J. G.; Adams, P. D.; Heazlewood, J. L.; Petzold, C. J. Standard Flow Liquid Chromatography for Shotgun Proteomics in Bioenergy Research. Front. Bioeng. Biotechnol. 2015. https://doi.org/10.3389/fbioe.2015.00044.

(19) Turpen, P. B.; Hockberger, P. E.; Meyn, S. M.; Nicklin, C.; Tabarini, D.; Auger, J. A. Metrics for Success: Strategies for Enabling Core Facility Performance and Assessing Outcomes. J. Biomol. Tech. 2016. https://doi.org/10.7171/jbt.16-2701-001.

(20) Kelstrup, C. D.; Bekker-Jensen, D. B.; Arrey, T. N.; Hogrebe, A.; Harder, A.; Olsen, J. V. Performance Evaluation of the Q Exactive HF-X for Shotgun Proteomics. J. Proteome Res. 2018. https://doi.org/10.1021/acs.jproteome.7b00602.

(21) Bache, N.; Geyer, P. E.; Bekker-Jensen, D. B.; Hoerning, O.; Falkenby, L.; Treit, P. V.; Doll, S.; Paron, I.; Müller, J. B.; Meier, F.; et al. A Novel LC System Embeds Analytes in Pre-Formed Gradients for Rapid, Ultra-Robust Proteomics. Mol. Cell. Proteomics 2018. https://doi.org/10.1074/mcp.TIR118.000853.

(22) Shen, Y.; Van Beek, T. A.; Zuilhof, H.; Chen, B. Hyphenation of Optimized Microfluidic Sample Preparation with Nano Liquid Chromatography for Faster and Greener Alkaloid Analysis. Anal. Chim. Acta 2013. https://doi.org/10.1016/j.aca.2013.08.034.

(23) Shi, J.; Wu, F.-X. Protein Inference by Assembling Peptides Identified from Tandem Mass Spectra. Curr. Bioinform. 2009. https://doi.org/10.2174/157489309789071048.

